# Analysis of aroma compounds in Barbecued Mutton (Kebab) during different storage periods and exploration of customer preferences

**DOI:** 10.1101/2025.02.16.638556

**Authors:** Aygul Alim

**Affiliations:** School of Life Science, Zhuhai College of Science and Technology, Jinwan, China

**Keywords:** Kebab, Storage period, GC-IMS, Aroma components, Molecular docking

## Abstract

Kebab (barbecued mutton) is a traditional meat product with a rich roasted meaty aroma and is highly favored by consumers. However, there are no reports on the flavor changes of different roasted meat tissues during storage, and there are no studies on why consumers prefer to eat Kebab. This study employed GC-IMS to explore the changes in aroma compounds in Kebab. Simultaneously, molecular docking was used to explore the mechanism of aroma compounds that trigger pleasure and to infer the optimal storage methods. A total of 61 volatile compounds, including aldehydes, alcohols, esters, ketones, acids, furans, ethers, and terpenes, were identified. Sensory evaluation revealed that when the storage time increased, the sour and sulfur aromas of mutton were strong, whereas the fatty aroma significantly increased and decreased in fat and lean meat. Additionally, a sense of pleasure was speculated by exemplifying six key aroma compounds. The differences in aroma compounds in the Kebab samples were clearly observed and indicated that the lean meat kebab maintained numerous meaty aroma compounds during storage until the 28th day. This is the first study to reveal the potential mechanism of the pleasure generated by eating Kebab, which will provide practical application value for evaluating the flavor of Kebab.

## INTRODUCTION

Xinjiang is one of the five major animal husbandry bases in China. Owing to its unique natural environment, the mutton produced here has the characteristics of tender meat, a delicious taste, and no muttony smell or greasiness. It is deeply loved by consumers and has been the main source of meat consumed by the people of Xinjiang for a long period of time. Barbecued meat products, with their strong aroma and unique cultural attributes, are deeply loved by customers. As a traditional meat product of ethnic groups such as the Uyghur people, Kebab (barbecued mutton) has unique characteristics in color, aroma, taste, and shape. Clients often evaluate product quality on the basis of these characteristics. The level of lipid oxidation in meat is closely linked to its edibility and storage characteristics and is also a major prerequisite for the formation of meat flavor^1^. Compared with other meat products, Kebab is a nutritious and healthy food with high protein, low fat, and low cholesterol contents. It is equally easy to make, delicious, and affordable. Kebab can be used as a street-style fast food or a delicious dish for entertaining guests. It meets the life requirements of people in this modern age. Given the enormous consumer market demand, relevant research on barbecued mutton is extremely urgent. However, owing to the scarcity of packaged barbecued mutton products and the lack of unified relevant parameters, it is difficult to carry out innovative industrial production.

Gas chromatography-ion mobility spectrometry (GC-IMS) is an emerging gas separation and detection technology that combines the high separation capacity of gas chromatography with the rapid response feature of ion mobility spectrometry (IMS). This combination endows GC-IMS with rapid analysis, high sensitivity, and variable-volume injection ability, all without the need for pretreatment^2^. In recent years, GC-IMS has developed rapidly in food flavor analysis and has been used in various food research fields, including establishing fingerprint spectra of volatile flavor compounds for food classification and adulteration detection^3,4^, investigating the changes in volatile substances during meat processing and volatile components during storage^5,6,7,8,9,10^, evaluating the freshness and degree of spoilage of food^11^, and detecting food odors^12,13^. However, GC-IMS technology has not been used to characterize the fingerprint of volatile flavor substances in different tissues of Kebab during storage.

During food processing or storage, odorless food ingredients can be transformed into odor-active volatiles through biochemical or chemical reactions. The knowledge about these precursors and reactions that result in the formation of odor substances in the food manufacturing process can provide the possibility of enhancing product flavor or preventing odors by selecting appropriate raw materials or optimizing processing conditions. On the other hand, Kebab has traditionally been favorite among consumers because people’s senses are delighted by the delicious aroma and taste of Kebab. In recent years, computer-based molecular modeling techniques such as molecular docking and molecular dynamics have received widespread attention because of their low cost and high time efficiency. In the context of odor-induced taste enhancement (OITE) and sugar reduction, these techniques have been used to explore odorants, and it has been indicated that beef odors potentially lead to a sense of pleasure^14^.

This study used GC-IMS technology to collect samples of barbecued mutton (Kebab) once a week during low-temperature refrigeration for a total of 28 days, with 5 tests for volatile organic compounds. These findings lay the groundwork for further research and development of the freshness and flavor components of Kebab. Furthermore, it can enhance the flavor of lamb, remove odors, and produce processed mutton products with excellent flavors to meet increasing demands for flavor and quality. On this basis, spectroscopic methods and molecular simulation techniques were used to study the mechanism of pleasure in response to the Kebab flavor and combined with sensory experiments to explore the physiological mechanism of pleasure caused by Kebab, providing a theoretical basis for supplementing nutritional and spiritual pleasure with roasted mutton products.

## MATERIALS AND METHODS

### Materials and Instruments

The newly barbecued mutton was prepared by a very famous Barbecue (Kebab) restaurant in Urumqi city, China. The lab instruments for the analysis were a Flavor Spec ® gas phase ion mobility spectrometer manufactured by G.A.S. Gesellschaft für analytische Sensorsysteme mbH, Germany.

### Methods

#### Sample preparation for experiments

The ordinary method of roasting mutton involves randomly selecting deboned mutton, mixing fat and lean meat, adding salt and spices (crystalline powder, chili powder) and then roasting. The purpose of this study was to explore flavor changes and the sources of flavors during the storage of mutton kebabs. Therefore, the roasted mutton in this experiment was divided into lean meat only, fat meat only, and fat-lean mixed meat (KL, KF, and KM). A total of 500 g of sample was taken and divided into four parts. Each divided barbecued mutton sample was divided into three parallel parts, and 4.5 g (accurate to 0.01 g) of each parallel sample was weighed and then vacuum-packed in aluminum foil bags and stored at low temperature (0–4°C) for later use. Samples were taken once a week. Three grams of each sample were accurately weighed, placed in a 20 mL headspace vial, and sealed. The parameters of the automatic sampler were as follows: the incubation temperature of the headspace vial was 60°C, the incubation time was 15 minutes, the incubation speed was 500 r/min, the injection needle temperature was 85°C, and the injection volume was 500 μL. Three replicates were set for each sample.

The **GC-IMS conditions** were as follows column type: FS-SE-54-CB-1 (15 m × 0.53 mm, 1 μM), the column temperature was 60°C; automatic headspace injection: the injection volume was 500 μL, the incubation time was 20 min, the incubation temperature was 65°C, the injection needle temperature was 85°C, the incubation speed was 500 r/min, the carrier/drift gas was N2, and the analysis time was 30 min. The flow rate was initially 2 mL/min and was maintained for 2 min; then, the flow rate linearly increased to 15 mL/min within 10 min, increased to 100 mL/min within 20 min, and then increased to 150 mL/min within 30 min. Then, the data processing began.

#### Sensory evaluation

Sensory evaluation was performed in three different sessions inside a sensory evaluation room at 20–24°C. The sensory evaluation team had 12 trained panellists (aged 23–40 years; 50% female and 50% male; 5 teachers and 7 graduate students at our College of Life Sciences; this demographic information was added as a covariate in the statistical analysis). On the evaluation day, the samples for all the sessions were placed under the same conditions. A total of 12 panelists were present in each session, with 3 rounds of testing. All the panelists had two weeks of sensory evaluation training. The characteristics of the samples were scored via a 10-point system. The intensities of the aroma and taste were ranked on a 10-point scale (0–2, very weak; 3–4, weak; 5–6, medium; 7–8, strong; 9–10, very strong; ^15^. The experiments were performed in triplicate, and the average scores of each trait were used for the statistical analysis.

#### Statistical analyses

The software FlavorSpec® fragrance analyzer (VOCal, Gallery Plot, Reporter, DynamicPCA, etc.) was used for the analysis. The statistics and analysis of the data were conducted via Office Excel 2007. The multimodal model Deep-B3 tool was used to predict flavor compounds.

## RESULTS

### Comparison of flavor compounds in different samples via GC-IMS

The Reporter plugin of GC-IMS was used to generate GC-IMS three-dimensional maps of volatile compounds in different tissues of Kebab (barbecued mutton) samples during storage. The three-dimensional map of the GC-IMS data shown in Figure 1 includes the retention time, transfer time, and peak intensity. In Fig. 1, each coordinate represents a different retention time, migration time, and peak intensity, and each peak signal represents a volatile organic compound, while the three-dimensional image displays the presence of volatile organic compounds. As shown in Fig. 1, by measuring the residual time and migration time in a sample, different volatile components can be identified, and peak intensities can be compared between different compounds. Two-dimensional GC-IMS spectra are more intuitive and easier to read, providing an accurate and comprehensive image of the characteristics and intensity of aromatic compounds, which is very useful for in-depth statistical analysis.

**Figure 1.**
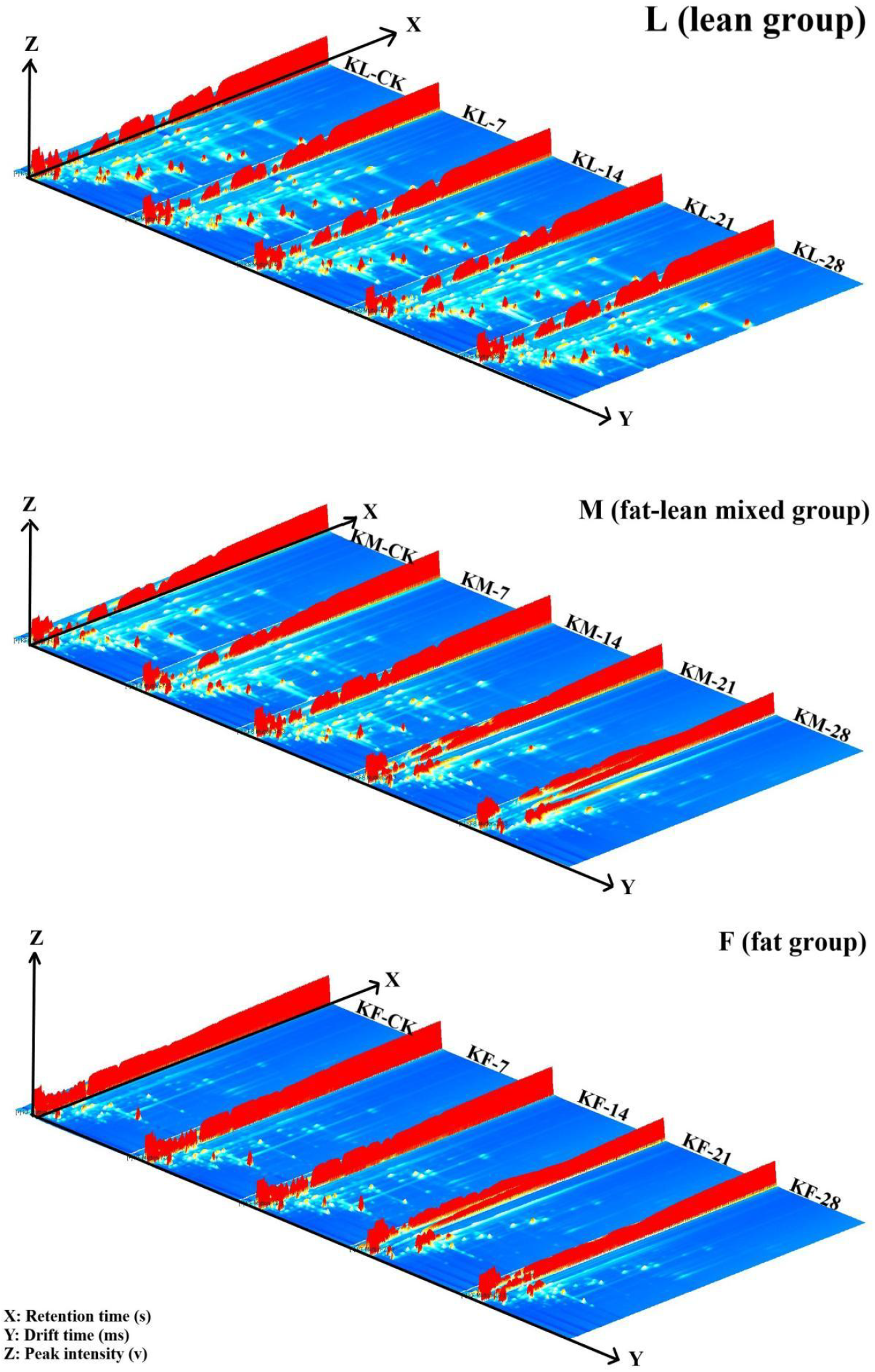
GC-IMS three-dimensional map of different tissues of Kebab samples during storage

Fig. 2 shows the GC-IMS spectra (vertical view) of different tissues from the Kebab samples at different storage times, including KL-CK (storage day 0), KL-7, KF-7, KFS (storage day 7), KL-14 (storage day 14), KL-21 (storage day 21), and KL-28 (storage day 28). The background of the entire graph is blue, with a horizontal axis of 1.0. The red vertical axis represents the RIP peak (the standardized reaction ion peak), the vertical axis represents the gas chromatography retention time, and the horizontal axis represents the ion migration time (standardized process). There are many points on both sides of the RIP peak, with one point for each volatile compound^16,17^. The colors of the two dots on both sides of the RIP peak represent the concentrations of red dots and light blue substances in the volatile organic roasted lamb sample. White indicates a lower substance content, red indicates a higher substance content, and darker peak colors indicate higher substance concentrations. Fig. 2 shows that the number and area of color spots on days 0 and 28 are significantly greater than those on days 7, 14, and 21 of storage, indicating a significant increase in the content of volatile substances on days 0, 28, and 0^18^. As shown in Fig. 2, there were significant differences in the volatile substance content of each sample. The red color of different tissues in lamb on day 0 was clearly greater than that on days 7, 14, 21, and 28, indicating that roasted lamb (stored on day 0) produced a large amount of volatile substances. The conclusion is similar to that shown in Fig. 2. To clearly compare different samples, the following are the fingerprint spectra of each peak^19^.

**Figure 2.**
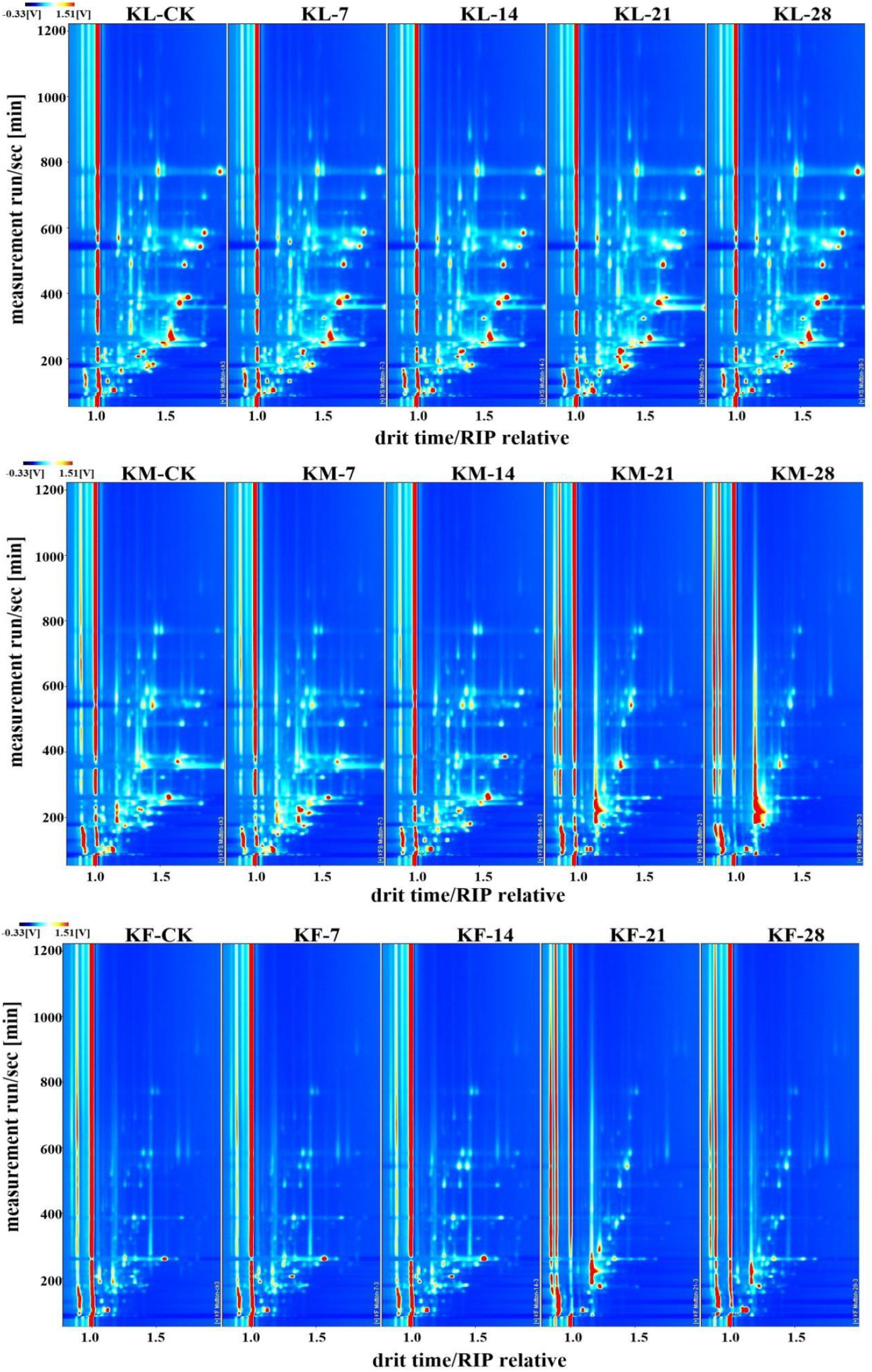
Comparison of HS-GC-IMS two-dimensional spectra of different tissues of Kebab samples during storage (vertical view)

As shown in Fig. 3, through the fingerprint spectrum, the commonalities and differences in the composition and content of flavor substances among the three groups of kebab tissues during the storage period can be visually compared. In Fig. 3, each row represents the composition of volatile components detected in one sample, and each column represents the signal peak situation of the same volatile flavor component in different samples. The fingerprint spectrum is a comparison of volatile substances in three types of mutton during five stages within 28 days of storage. The three groups of samples and 15 different samples can be distinguished by volatile substances. Moreover, most of the volatile substances in lean meat (KL) have relatively high concentrations, and the concentrations of volatile substances during the entire storage period are higher than those in fat meat (KF) and the mixture of lean and fat meat (KM). A separate discussion of each group revealed that the volatile substances in each group of samples varied with different storage periods. In the KF group, KF-CK to KF-14 formed one group, and KF-21 to KF-28 formed another group. Among the three groups, the KF group clearly contained the fewest types of volatile substances. In the figure, the substances in area A have the highest concentration in KL, followed by those in KM, and the lowest or no substances are present in KF. The pattern of substances in this area may be related to the content of lean meat (KL). The substances in this area from left to right mainly include β-pinene, γ-terpinene, butyric acid acetate, cuminaldehyde, E-2-nonanal, E-2-octanal, 2-pentylfuran, 2-hexanal, nonanol, nonanal, 3-methylbutanal, 2-pentanone, hexanol, E-2-heptanal, hexyl acetate, heptanal, E-2-pentanal, butanal, butyl 2-methylbutyrate, acetoin, and octanal. In the figure, the substances in area B have relatively high concentrations during the storage period in KM, relatively high concentrations in the later storage period of KF, and relatively low or none during the entire storage period of KS. They mainly include 2-octanol, 1-hydroxy-2-propanone, ammonia, trimethylamine, triethylamine 2-methylbutanal, and 2-methyl-1-propanol. In the KM group, the volatile substances of the KM-14 sample clearly do not conform to the overall pattern. These results indicate that the volatile components of different tissues of roasted meat are different and that the types and contents of volatile components also change with storage time^18^.

**Figure 3.**
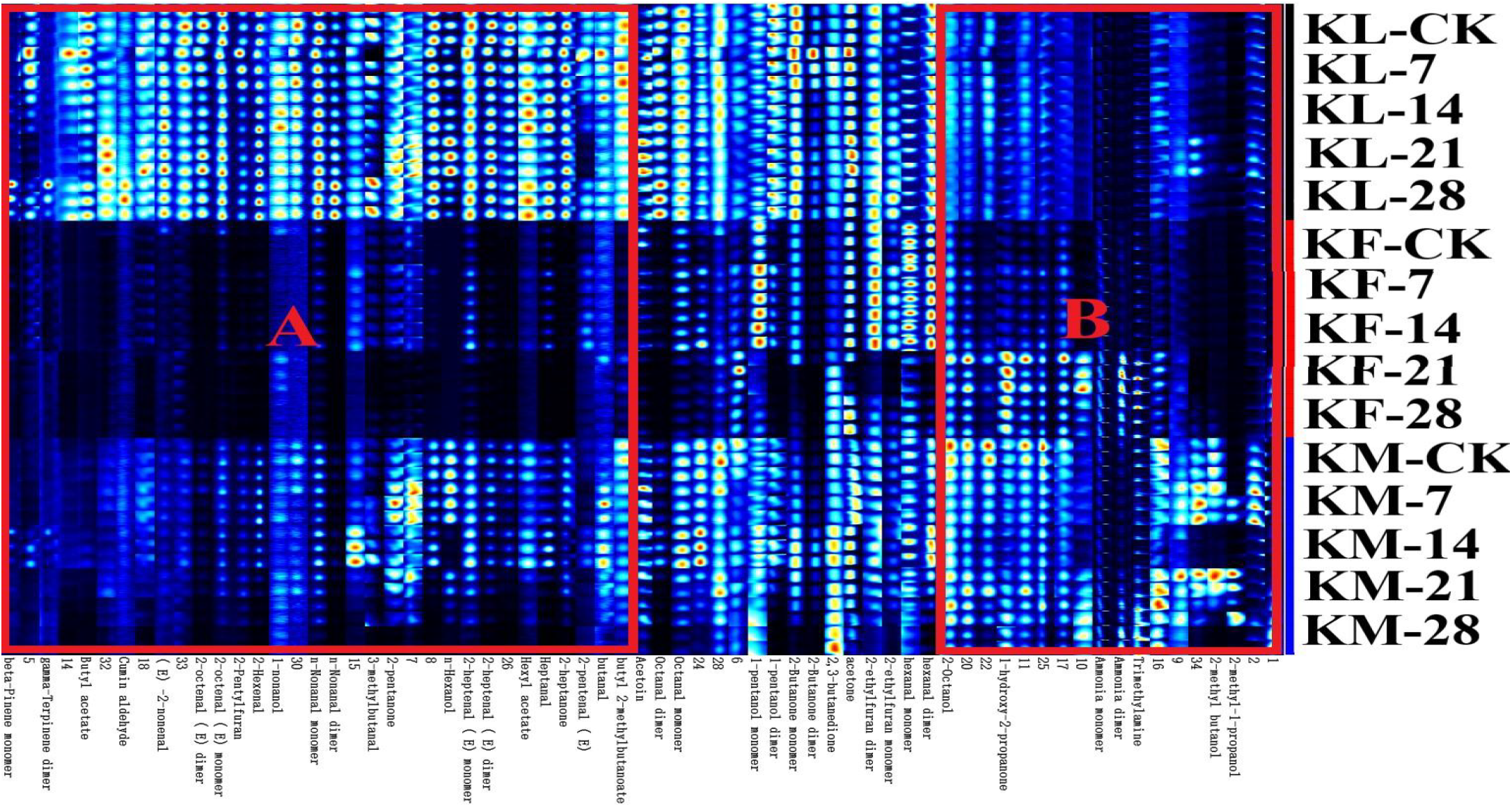
Gallery plot (fingerprint map)

The fingerprint spectrum is a comparison of the volatile substances of the mutton in the KL group. There were significant differences in the quantities and concentrations of the volatile compounds among the five samples. The figure shows that there are different compounds in different storage periods; that is, some substances have the highest concentrations in specific stages and lower concentrations or are absent in other stages. For example, the aroma compounds in area A in the figure have the highest concentration in KL-CK. The compounds with the highest concentrations in KL-7 mainly include 2-butanone, 5-methyl-2-furancarbaldehyde, E-2-pentanal, 2-pentylfuran, butyl acetate, butyl 2-methylbutyrate, 2-heptanone, 2,3-butanedione, etc. The substances with relatively high concentrations in KL-14 mainly include nonanol, etc. The substances with the highest concentrations in KL-21 mainly include (E)2-nonanal, 2-pentanone, 1-hydroxy-2-propanone, (E)2-octanal and hexanol, etc. The substances with the highest concentrations in KL-28 mainly include 3-methylbutanal, β-pinene, γ-terpinene, 2-hexanal and nonanal, etc.

In the KF group, KF-CK to KF-14 formed one group, and KF-21 to KF-28 formed another group. The concentration in KF-CK was low, and the concentrations in KF-7 and KF-14 were the same and greater than those in KF-21 and KF-28. The volatile substances from left to right mainly include 2-butanone, 3-methylbutanal, 2-pentanone, 2-pentylfuran, octanal, hexyl acetate, nonanal, (E)2-octanal, heptanal, 2-acetylfuran, pentanol, hexanal, ethyl hexanoate, (E)2-heptanal, (E)2-pentanal, and 2-hexanal, etc. The concentrations of the following compounds are the same in KF-21 and KF-28 and higher than those in the other samples, mainly trimethylamine, butyl 2-methylbutyrate, hydroxy-2-propanone, and ammonia, etc. In terms of fat oxidation, the concentrations of these compounds increased with increasing storage time.

In the KM group, it is obvious that the KM-14 sample has a relatively large difference from the others. The compound concentrations were relatively high in KM-7 and KM-21. From left to right, the main copounds were 2-methylbutanal, 2-pentanone, 2-hexanal, acetoin, butanal, octanal, pentanol, acetone, 2-butanone, 3-methylbutanal, butyl acetate, heptanal, (E)2-pentanal, β-pinene, nonanal, and hexyl acetate, etc. The compound concentration was the lowest in KM-28. The main substances were 1-octen-3-ol, 2-octanol, (E)2-octanal, (E)2-nonanal, butyl2-methylbutyrate, 2-heptanone, nonanol, hexanol, 2-pentylfuran, (E) 2-heptanal, 2-acetylfuran, etc. The concentration was the highest in KM-28. The main substances were 1-hydroxy-2-propanone, 2,3-butanedione, 2-methyl-1-propanol, trimethylamine, ammonia, etc. In terms of detecting the key aroma compounds of roasted mutton, 33 and 30 compounds were detected in mutton before and after roasting, respectively^11^. 3-Methylbutanal, pentanal, hexanal, heptanal, octanal, nonanal, and 1-octen-3-ol are important flavor compounds. Among them, hexanal and 1-octen-3-ol are key flavor compounds^20^. These results are similar to those of the present study, but there are no reports on the flavor changes of different tissues during the storage period.

### Qualitative analysis of aroma compounds in samples (GC×IMS Library Search)

To further analyze the changes in volatile compounds during storage in Kebab, qualitative analysis of substances can be performed via the database and IMS database. As shown in Fig. 4. The numbers in the figure indicate the characteristic peak position points of the volatile components of the samples. Each marked point represents a specific volatile component for qualitative analysis. The points on the entire spectrum represent all the volatile compounds detected in the sample^11^. There are 61 volatile substances that can be clearly qualitatively determined, including monomers and dimers of some volatile substances. The specific information is shown in Table 1.

**Table 1.** List of qualitatively determined compounds in GC-IMS (corresponding to the serial numbers in Figure 6).

| Count | Compound | CAS# | Formula | MW | RI | Rt [sec] | Dt [RIPrel] |
| --- | --- | --- | --- | --- | --- | --- | --- |
| 1 | Trimethylamine | C75503 | C3H9N | 59.1 | 736.7 | 215.082 | 1.16806 |
| 2 | hexanal monomer | C66251 | C6H12O | 100.2 | 791 | 260.48 | 1.26303 |
| 3 | hexanal dimer | C66251 | C6H12O | 100.2 | 791.4 | 260.925 | 1.56777 |
| 4 | 1-pentanol monomer | C71410 | C5H12O | 88.1 | 774.6 | 245.345 | 1.25226 |
| 5 | 1-pentanol dimer | C71410 | C5H12O | 88.1 | 773.5 | 244.455 | 1.50929 |
| 6 | Acetoin | C513860 | C4H8O2 | 88.1 | 747.2 | 223.489 | 1.33909 |
| 7 | n-Hexanol | C111273 | C6H14O | 102.2 | 878.4 | 356.72 | 1.99722 |
| 8 | 2-heptanone | C110430 | C7H14O | 114.2 | 893.3 | 373.866 | 1.6395 |
| 9 | Heptanal | C111717 | C7H14O | 114.2 | 901.4 | 388.399 | 1.70066 |
| 10 | n-Nonanal monomer | C124196 | C9H18O | 142.2 | 1108.9 | 776.21 | 1.46949 |
| 11 | n-Nonanal dimer | C124196 | C9H18O | 142.2 | 1107.2 | 772.918 | 1.95338 |
| 12 | 1,8-Cineole monomer | C470826 | C10H18O | 154.3 | 1043.2 | 648.093 | 1.29598 |
| 13 | 1,8-Cineole dimer | C470826 | C10H18O | 154.3 | 1039.6 | 641.241 | 1.73041 |
| 14 | 2-heptenal (E) monomer | C18829555 | C7H12O | 112.2 | 961.3 | 494.61 | 1.26241 |
| 15 | 2-heptenal (E) dimer | C18829555 | C7H12O | 112.2 | 958.2 | 489.129 | 1.67261 |
| 16 | 1-hydroxy-2-propanone | C116096 | C3H6O2 | 74.1 | 690.6 | 178.332 | 1.2327 |
| 17 | 2-Butanone monomer | C78933 | C4H8O | 72.1 | 593.3 | 135.077 | 1.07863 |
| 18 | 2-Butanone dimer | C78933 | C4H8O | 72.1 | 590.1 | 133.668 | 1.25106 |
| 19 | ethyl acetate | C141786 | C4H8O2 | 88.1 | 606.2 | 140.713 | 1.3459 |
| 20 | acetone | C67641 | C3H6O | 58.1 | 524.4 | 105.018 | 1.13893 |
| 21 | 2-pentanone | C107879 | C5H10O | 86.1 | 688.8 | 176.879 | 1.37869 |
| 22 | butanal | C123728 | C4H8O | 72.1 | 595.4 | 136.017 | 1.3007 |
| 23 | 2-methyl-1-propanol | C78831 | C4H10O | 74.1 | 634.2 | 152.925 | 1.3758 |
| 24 | 3-methylbutanol | C590863 | C5H10O | 86.1 | 651.4 | 160.44 | 1.4148 |
| 25 | Isopentyl alcohol | C123513 | C5H12O | 88.1 | 744.1 | 221.029 | 1.49135 |
| 26 | Triethylamine | C121448 | C6H15N | 101.2 | 707.7 | 191.909 | 1.46391 |
| 27 | 2-ethylfuran monomer | C3208160 | C6H8O | 96.1 | 727.7 | 207.878 | 1.04794 |
| 28 | 2-ethylfuran dimer | C3208160 | C6H8O | 96.1 | 727.7 | 207.878 | 1.31514 |
| 29 | 2-pentenal (E) | C1576870 | C5H8O | 84.1 | 751.2 | 226.642 | 1.10843 |
| 30 | ethyl 2-methylpropanoate | C97621 | C6H12O2 | 116.2 | 747.8 | 223.912 | 1.5742 |
| 31 | 2-methyl butanol | C137326 | C5H12O | 88.1 | 733.5 | 212.519 | 1.50588 |
| 32 | 2-Hexenal | C505577 | C6H10O | 98.1 | 848.9 | 324.331 | 1.52425 |
| 33 | Butyl acetate | C123864 | C6H12O2 | 116.2 | 818.5 | 290.833 | 1.61403 |
| 34 | alpha-Pinene monomer | C80568 | C10H16 | 136.2 | 925.3 | 430.71 | 1.22225 |
| 35 | alpha-Pinene dimer | C80568 | C10H16 | 136.2 | 922.7 | 426.068 | 1.67721 |
| 36 | beta-Pinene monomer | C127913 | C10H16 | 136.2 | 972.3 | 514.26 | 1.22842 |
| 37 | beta-Pinene dimer | C127913 | C10H16 | 136.2 | 972.9 | 515.292 | 1.64328 |
| 38 | beta-Pinene polymer | C127913 | C10H16 | 136.2 | 973.5 | 516.323 | 1.72965 |
| 39 | Octanal monomer | C124130 | C8H16O | 128.2 | 1011.4 | 586.297 | 1.40162 |
| 40 | Octanal dimer | C124130 | C8H16O | 128.2 | 1009.9 | 583.329 | 1.83775 |
| 41 | 5-Methyl-furfural | C620020 | C6H6O2 | 110.1 | 977.1 | 522.703 | 1.4787 |
| 42 | 1-octen-3-ol | C3391864 | C8H16O | 128.2 | 990.2 | 546.021 | 1.72872 |
| 43 | Ethyl hexanoate monomer | C123660 | C8H16O2 | 144.2 | 1005.6 | 574.85 | 1.33395 |
| 44 | Ethyl hexanoate dimer | C123660 | C8H16O2 | 144.2 | 1007.1 | 577.818 | 1.80767 |
| 45 | 2-Pentylfuran | C3777693 | C9H14O | 138.2 | 999.3 | 562.555 | 1.25687 |
| 46 | 2-Octanol | C123966 | C8H18O | 130.2 | 1010.4 | 584.177 | 1.4599 |
| 47 | Hexyl acetate | C142927 | C8H16O2 | 144.2 | 1010.6 | 584.601 | 1.89415 |
| 48 | gamma-Terpinene monomer | C99854 | C10H16 | 136.2 | 1063.7 | 688.046 | 1.22492 |
| 49 | gamma-Terpinene dimer | C99854 | C10H16 | 136.2 | 1063.4 | 687.481 | 1.70394 |
| 50 | 2-octenal (E) monomer | C2548870 | C8H14O | 126.2 | 1072.4 | 705.005 | 1.33771 |
| 51 | 2-octenal (E) dimer | C2548870 | C8H14O | 126.2 | 1068.5 | 697.495 | 1.82865 |
| 52 | 4-Terpineol | C562743 | C10H18O | 154.3 | 1177.7 | 910.284 | 1.22685 |
| 53 | (E) -2-nonenal | C18829566 | C9H16O | 140.2 | 1165.1 | 885.757 | 1.41477 |
| 54 | Cumin aldehyde | C122032 | C10H12O | 148.2 | 1259 | 1068.601 | 1.33543 |
| 55 | 1-nonanol | C143088 | C9H20O | 144.3 | 1165.7 | 886.827 | 1.53357 |
| 56 | Myrcene | C123353 | C10H16 | 136.2 | 990.9 | 547.161 | 1.29988 |
| 57 | Limonene | C138863 | C10H16 | 136.2 | 1034.2 | 630.577 | 1.65752 |
| 58 | butyl 2-methylbutanoate | C15706737 | C9H18O2 | 158.2 | 1042.2 | 646.302 | 1.37294 |
| 59 | 2,3-butanedione | C431038 | C4H6O2 | 86.1 | 587.6 | 132.59 | 1.16482 |
| 60 | Ammonia monomer | C7664417 | H3N | 17 | 903.3 | 391.754 | 0.88942 |
| 61 | Ammonia dimer | C7664417 | H3N | 17 | 901.3 | 388.096 | 0.84706 |

**Figure 4.**
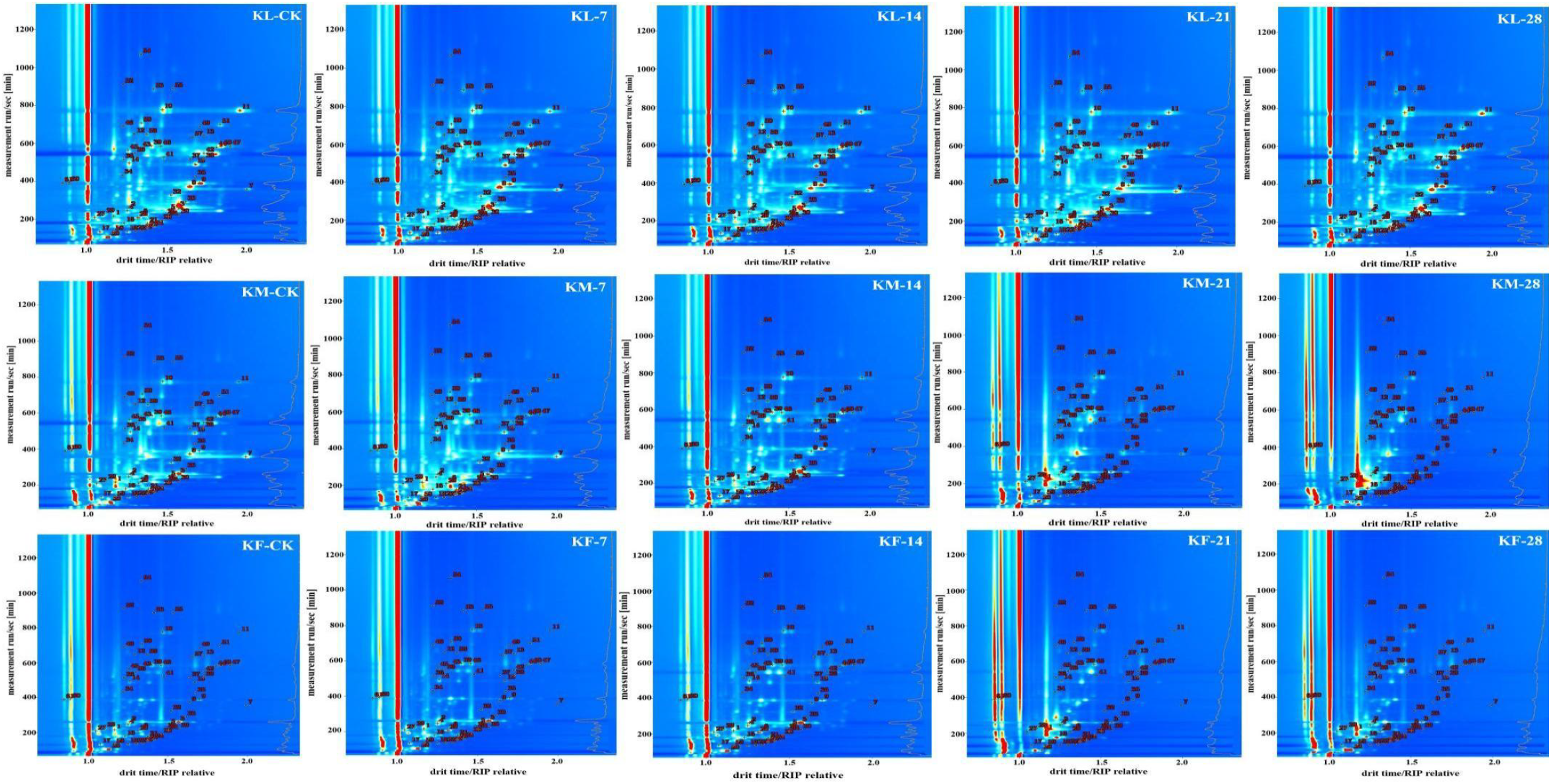
GC-IMS spectrum of the sample for qualitative analysis

As shown in Table 1, 61 volatile compounds were detected, including 13 aldehydes, 11 alcohols, 8 esters, 6 ketones, 1 acid, 2 furans, 1 ether, and 1 terpene. Aldehydes were the most abundant volatile components identified during the storage period of kebab, with a total of 13 volatile components. These include heptanal, butanal, 2-pentenal, 2-hexenal, and 5-methylfurfural. Second, the most abundant are alcohols, with a total of 11 kinds. These include 1-pentan-hexanol, 2-methyl-1-propanol, and 2-methylbutanol. Alcohol compounds mainly originate from the oxidation of unsaturated fats. Some alcohols may also be obtained by the reduction of aldehyde substances. Finally, the most abundant are esters, with a total of 8 kinds, such as ethyl acetate, ethyl2-methylpropanoate, butyl acetate, and hexyl acetate. Esters are important volatile compounds with relatively low thresholds but help improve the overall flavor quality of roasted mutton^11^. There are a total of 6 types of ketones, such as 2-heptanone and 1-hydroxy-2-propanone. Ketones are intermediates in the formation of heterocyclic compounds and have relatively high flavor thresholds. The least common compounds are acids, furans, and terpenes. However, both 2-ethylfuran and 2-pentylfuran promote the formation of plant aromatic flavors in meat products during the heating process. Wei et al. confirmed that hexal, octylaldehyde, 1-octene-3-ol, aldehyde, heptal, pentalaldehyde, 3-methylbutyraldehyde, and ethylfuran are key aroma compounds in roasted mutton^20^.

### PCA diagram of samples

Consistent with the Gallery plot shown in Fig. 5, PCA can distinguish the five roasted mutton samples. If the samples are close to each other, there is a small difference. If they are far apart^21,22^.

**Figure 5.**
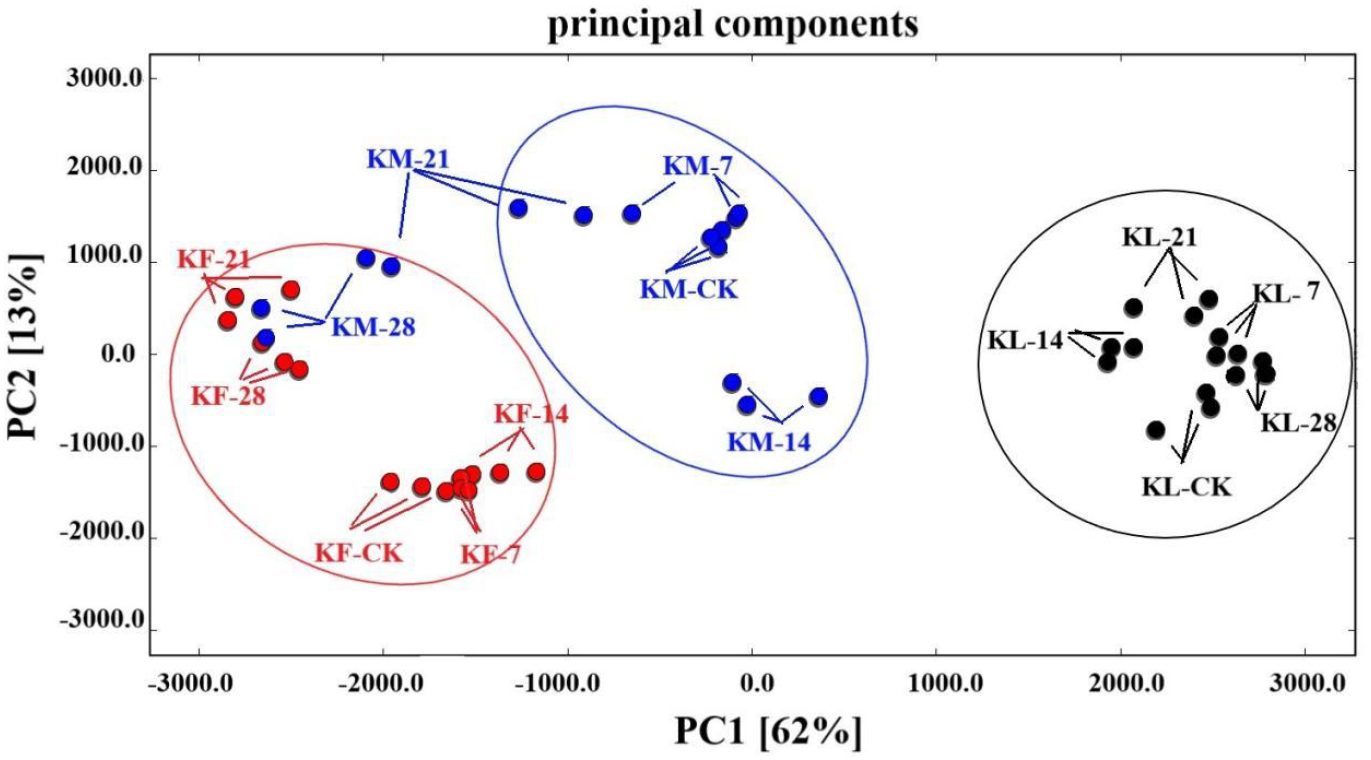
PCA diagram of samples

The samples of KL, KF and KM within 28 days were subjected to cluster analysis via the PCA plug-in. The results are shown in Fig. 5. The results show that when GC-IMS and principal component analysis are used, the kebab at different refrigeration times can be identified. The variation contribution rates of PC1 and PC2 are 13% and 62%, respectively, and the cumulative variance contribution reaches 74%. For the first principal component (PC1), the distances between KF and KM are close, indicating that they have high similarity in volatile substance types and similar flavors. The distance on the 21st day was greater than that at the other times, indicating that there were obvious differences in the types and flavors of the volatile compounds. For the second principal component (PC2), the distances on the 7th, 14th, and 28th days of the KM samples are close, indicating that there is high similarity in volatile substance types and flavors and that the distance on the KM-21 samples is still greater than that at other times. Intuitively, the distance between the KL group and KM is very close, indicating that they have similar volatile compounds. This may contribute to the similarity in their volatile compound compositions.

### Sensory evaluation of the aroma

Fig. 6A (lean), B (mixed meat), and C (fat) show the aroma analysis results for Kebab stored for different durations. Sensory evaluation revealed that meat, nutty, fatty, green, sulfur, and sour aromas were the predominant aromas of the samples. In samples A and B, meat had the strongest aroma, and in sample C, the fatty aroma was the most predominant. Among all the samples, the sour aroma was the weakest. When the storage time reached 28 days, sample A maintained strong meaty and nutty aromas; in samples B and C, the fatty, sulfur, and sour aromas increased significantly. The results show that the sour and sulfur aromas in mutton became stronger when the storage time increased. While the fatty aroma became more evident in the fat meat when the storage time increased, its intensity was significantly weaker in the lean meat.

**Figure 6.**
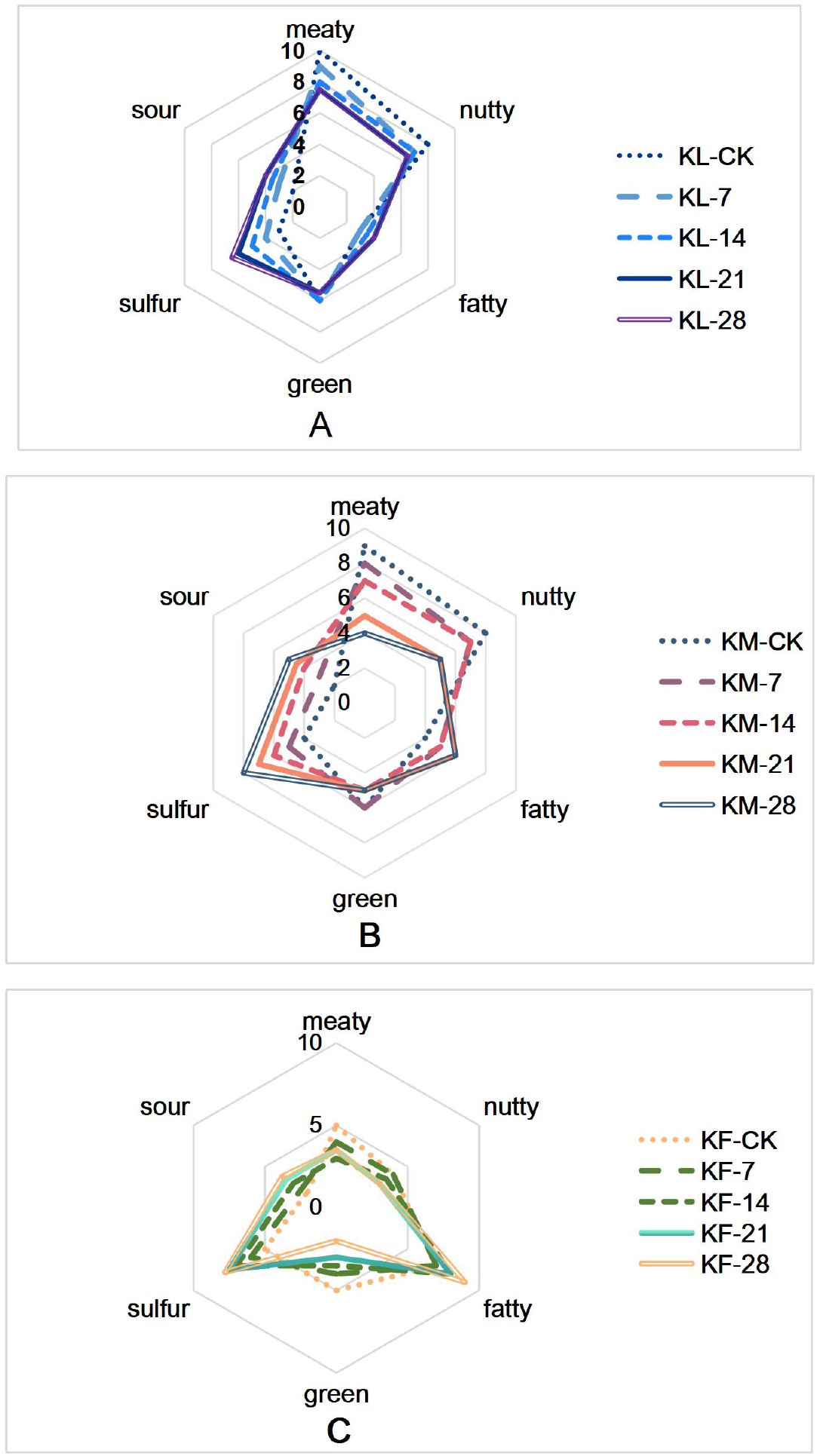
Aroma sensory evaluation results.

### Interaction between flavor substances that produce a sense of pleasure and serotonin (5-HT) receptors

#### Blood−brain barrier permeability prediction

The blood−brain barrier is a physiological barrier formed by vascular endothelial cells that limits the ability of many substances (such as most drugs) to enter brain tissue from the blood. Drugs that affect the central nervous system, including drugs that act on 5-HT receptors, usually need to be able to cross the blood−brain barrier to have an effect. This study revealed that these key flavor substances can make people feel pleasure; therefore, predicting whether they have blood−brain barrier penetration is a key indicator for determining their impact on the central nervous system. For this purpose, we used the deep learning-based multimodal model Deep-B3^23^ tool to predict six flavor compounds, namely, 5-methylfurfural, 2-pentylfuran, (E)-2-hexenal, 2-octanone, trans-2-nonanal, and 3-methylbutanol. The results in Table 2 show that all the samples penetrated the blood−brain barrier.

**Table 2.**
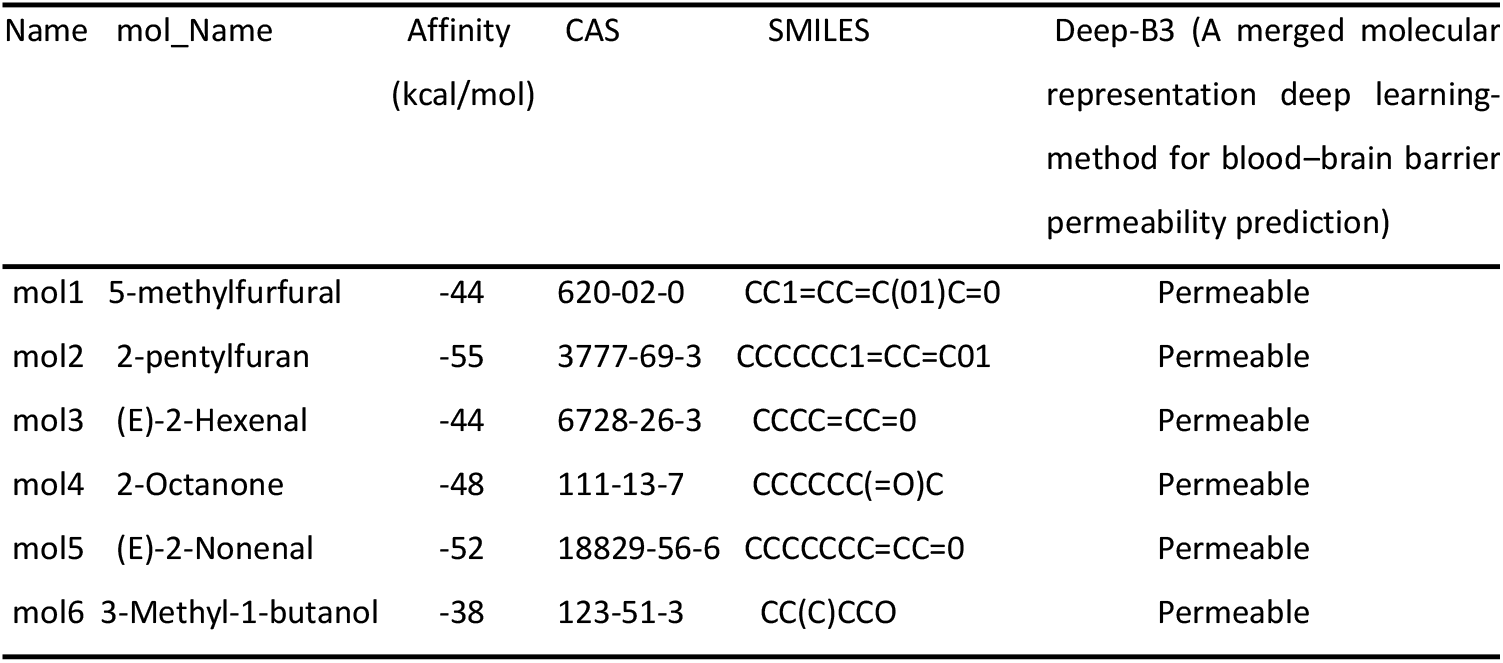
The deep learning-based multimodal model Deep-B3.

#### Molecular docking

The 5-HT1E receptor structure (PDB ID: 6G79) containing the binding-selective small-molecule agonist was downloaded from the RCSB database. The structures of the flavor substances were downloaded from the PubChem database. The MMFF94s force field is subsequently used for geometric optimization to eliminate unreasonable bond lengths and bond angles. Semiflexible molecular docking was carried out via the AutoDock Vina program^24^, and the cocrystallized small-molecule agonist binding region was used as the conformational search space (center_x = 94.3, center_y = 52.2, center_z = 64.3) to explore potential geometric sites. Afterwards, the best complex structure is selected according to the binding energy and binding conformation, and the Open Source PyMoL program is used for visual interaction analysis (Fig. 7).

**Figure 7.**
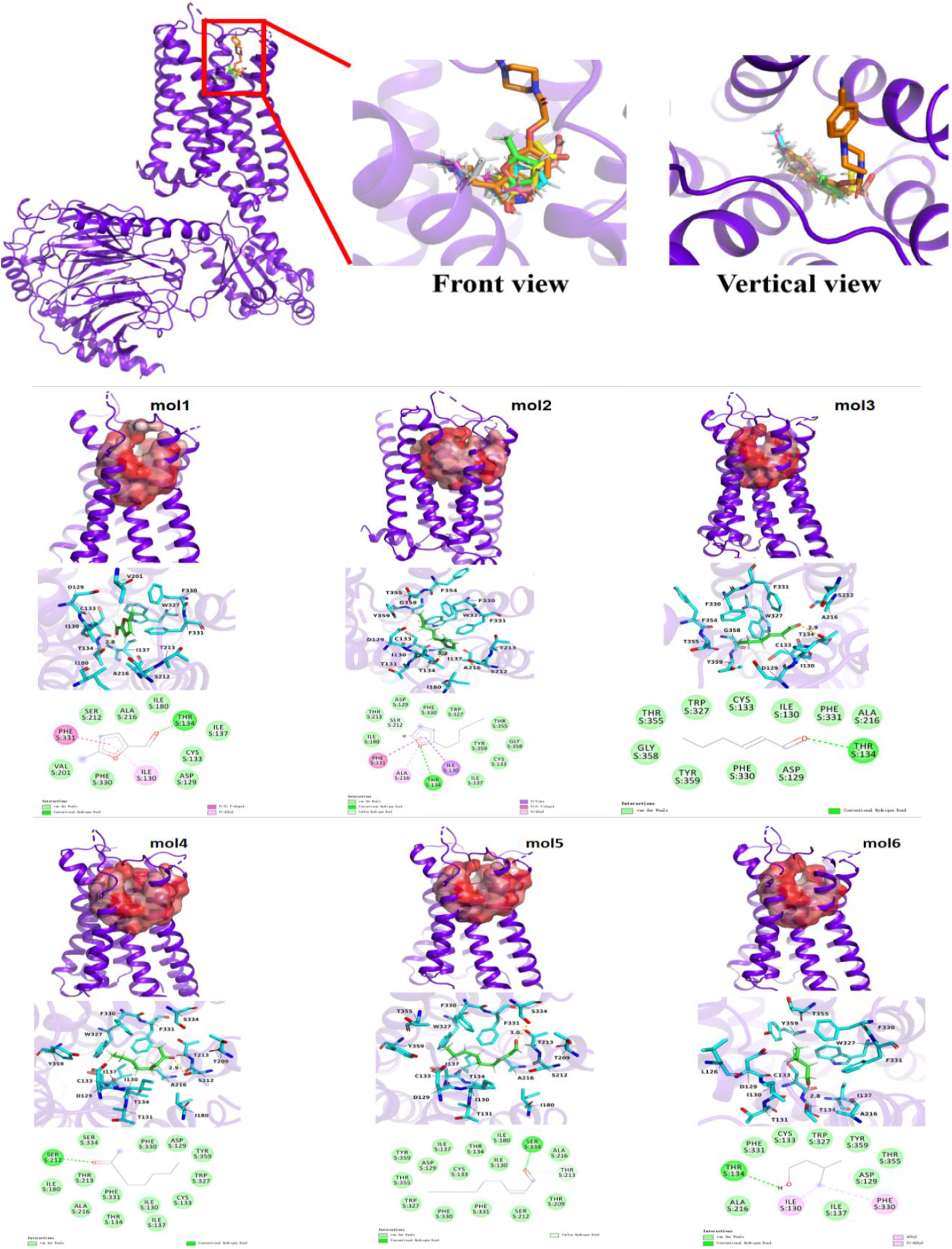
Comparison of the binding conformations of the six flavor molecules and the cocrystallized activator molecule

## DISCUSSION

In this study, the volatile components of kebab during cold storage were analyzed via GC-IMS. A total of 61 volatile substances were detected and identified, and the volatile compounds included aldehydes (acetaldehyde, N-nonanal, 2-octenal heptanal, butanal, 2-pentenal, 2-hexenal, 5-methylfurfural), alcohols (1-pentanol, n-hexanol, 2-methyl-1-propanol, 3-methylbutanoI, 2-methylbutanol, 1-octen-3-ol), esters (ethyl acetate, ethyl 2-methylpropanoate, butyl acetate, hexyl acetate, butyl 2-methylbutyrate), ketones (2-heptanone, 1-hydroxy-2-propanone, 2-butanone and acetone), furans (2-ethylfuran, 2-pentylfuran), ethers, terpenes, etc.

The results revealed that there are more key meaty aroma compounds in lean meat (KL) in terms of quantity and content. Although its shelf-life has increased, it still maintains a better roasted meat flavor. However, after the KM sample was stored for 21 days, the aroma components were similar to those of the KF sample. Even at 28 days, the flavor was very similar to that of the KF sample. Compounds with unpleasant aromas are produced, resulting in off-flavors. Fingerprint spectra were used to analyze the volatile components in different storage periods to determine the differences in volatile components between samples, and the results were the same as those mentioned above. The KM and KF samples contained fat tissue, which eventually led to similarity. Finally, the molecular mechanism of the production of a sense of pleasure by six key aroma compounds in kebab was verified via molecular docking simulation methods.

## CONCLUSIONS

At present, mutten production is mostly in the form of small workshops, with a small scale and inconsistent quality. kebab has soft and tender meat and a unique flavor. To achieve flavor fidelity, it is very important to study the characteristic qualities of traditional kebabs. During the storage process, the original quality of the lean meat tissue of roasted mutton is better than that of mixed meat and fatty meat. The aroma compounds of kebabs include mainly aldehydes, ketones, furans, alcohols and esters. Aroma deeply affects our emotions, memories and overall well-being. Eating kebabs to improve mood and produce a sense of pleasure is no exception. Studies have shown that in addition to the impact of aroma on emotions, fragrance may also have physiological effects and help reduce feelings of anxiety and depression. Therefore, further research on the impact of aroma on emotions is highly important.

## ACKNOWLEDGEMENTS

This research was funded by the ‘Three Levels’ Talent Construction Project of Zhuhai College of Science and Technology.

## DATA AVAILABILITY STATEMENT

The data that have been used are confidential.

## SUPPORTING INFORMATION

Supporting information may be found in the online version of this article.

## REFERENCES

1 Alim A, Song H, Liu Y, Zou, T, Zhang Y, Zhang S and Raza A, Research of beef-meaty aroma compounds from yeast extract using carbon module labeling (CAMOLA) technique. Lwt 112:108–239 (2019).

2 Wang S, Chen H and Sun B, Recent progress in food flavor analysis using gas chromatography– ion mobility spectrometry (GC–IMS). Food Chem 315:126–158(2019).

3 Othman A, Goggin KA, Tahir NI, et al., Use of headspace–gas chromatography-ion mobility spectrometry to detect volatile fingerprints of palm fiber oil and sludge palm oil in samples of crude palm oil. BMC Research Notes, 12:1–6 (2019).

4 Rodríguez-Maecker R, Vyhmeister E, Meisen S, et al., Identification of terpenes and essential oils by means of static headspace gas chromatography-ion mobility spectrometry. Anal and Bioanal Chem 409:6595–6603 (2017).

5 Yu Y, Wang G, Luo Y, Pu Y, Ge C, and Liao G, Effect of natural spices on precursor substances and volatile flavor compounds of boiled Wuding chicken during processing. Flav Fragr J 35(5):570–583.(2020)

6 Li H, Liu J, Wang Z, Liu X, Yan X, Liu S and He, X, Process optimization of chili flavor beef tallow and analysis of its volatile compounds by GC-IMS. Int J Food Engin 17(7):507–516 (2021).

7 Li W, Chen YP, Blank I, Li F, Li C and Liu Y, GC× GC-ToF-MS and GC-IMS based volatile profile characterization of the Chinese dry-cured hams from different regions. Food Res Int 142:110222 (2021).

8 Tian XY, Aheto JH, Huang X, Zheng K, Dai C, Wang C and Bai JW, An evaluation of biochemical, structural and volatile changes of dry-cured pork using a combined ion mobility spectrometry, hyperspectral and confocal imaging approach. J Sci Food Agri 101(14): 5972-5983 (2021).

9 Bi J, Lin Z, Li Y, Chen F, Liu S and Li C, Effects of different cooking methods on volatile flavor compounds of chicken breast. J Food Biochem 45(8):e13770 (2021).

10 Wen R, Kong B, Yin X, Zhang H and Chen Q, Characterization of flavor profile of beef jerky inoculated with different autochthonous lactic acid bacteria using electronic nose and gas chromatography–ion mobility spectrometry. Meat Sci 183:108658 (2022).

11 Liu H, Hui T, Fang F, Li S, Wang Z and Zhang D, The formation of key aroma compounds in roasted mutton during the traditional charcoal process. Meat Sci 184:108689 (2022).

12 Liu H, Hui T, Fang F, et al., Characterization and discrimination of key aroma compounds in preand postrigor roasted mutton by GC-O-MS, GC E-Nose and aroma recombination experiments. Foods 10(10):2387 (2021).

13 Pei F, Yang W, Ma N. et al., Effect of the two drying approaches on the volatile profifiles of button mushroom (Agaricus bisporus) by headspace GC–MS and electronic nose. LWT 72:343–350 (2016).

14 Jayasena DD, Jung S, Kim HJ, Alahakoon AU, Nam KC and Jo C, Effect of sex on flavor-related and functional compounds in freeze-dried broth made from Korean native chicken. Korean J Food Sci Anim Res 34:448–456 (2014).

15 Alim A, Song L,Yang C, Ye L and Zhou TT, Flavor-active compounds in thermally treated yeast extracts[J]. J Sci Food Agr 98(10):3774–3783 (2018).

16 Sun X, Yu Y, Wang Z, et al., Insights into flavor formation of braized chicken: Based on E-nose, GC–MS, GC-IMS, and UPLC-Q Exactive-MS/MS. Food Chem 448:138972 (2024).

17 Chen M, Chen T, Qi X, Lu D and Chen B, Analyzing changes of volatile components in dried pork slice by gas chromatography-ion mobility spectroscopy. CyTA-J Food 18(1):328–335 (2020).

18 Alim A and Hu WZ, A study with sensory evaluation and GC‒MS/HPLC analysis on changes of flavor compounds in mutton during storage. CyTA-J Food 22:2326060 (2024).

19 Ge S, Chen Y, Ding S, et al., Changes in volatile flavor compounds of peppers during hot air drying process based on headspace-gas chromatography-ion mobi lity spectrometry (HS-GC-IMS). J Sci Food Agr 100(7):3087–3098 (2020).

20 Wei J, Wang L, Ma X, Xu Z and Wang Z, Effects of Variable-Temperature Roasting on the Flavor Compounds of Xinjiang Tannur-Roasted Mutton. Foods 13(19):3077(2024).

21 Li MQ, Yang RW, Zhang H, Wang SL, Chen D and Lin SY, Development of a flavor fingerprint by HS-GC-IMS with PCA for volatile compounds of tricholoma matsutake singer. Food Chem, 290:32–39 (2019).

22 Wan J, Liu Q, Ma C, et al., Characteristic flavor fingerprint disclosure of dzo beef in Tibet by applying SAFE-GC-O-MS and HS-GC-IMS technology. Food Res Int 166:112581 (2023).

23 Tang Q, Nie F, Zhao Q and Chen W, A merged molecular representation deep learning method for blood–brain barrier permeability prediction. Brief in Bioinf 23(5):bbac357 (2022).

24 Trott O and Olson AJ, AutoDock Vina: improving the speed and accuracy of docking with a new scoring function, efficient optimization, and multithreading. J comp chem 31:455–461 (2010).

